# Fundamental limitations of genomic language models for realistic sequence generation

**DOI:** 10.64898/2026.01.17.700093

**Authors:** Alexandros Tzanakakis, Ioannis Mouratidis, Ilias Georgakopoulos-Soares

**Author notes:** These authors jointly supervised the work (Ioannis Mouratidis, Ilias Georgakopoulos-Soares).

## Abstract

Large language models (LLMs) have shown remarkable success in natural language processing, prompting interest in their application to genomic sequence analysis. Genomic Language Models (gLMs) based on similar architectures offer a promising avenue for synthetic genome generation and characterization. However, their effectiveness for biological sequence modeling remains poorly characterized. We present a comprehensive evaluation of genomic language models that explicitly aim to generate entire synthetic genomes. We tested Evo 2 on diverse prokaryotic, eukaryotic and viral genomes, and megaDNA on bacteriophage genomes, and assessed performance across key biological features and organizational patterns. Our results reveal systematic failures in gLM-based genomic reconstruction. While the synthetic sequences captured local sequence statistics, they consistently failed to preserve long-range genomic organization, repeat and k-mer composition, transcription factor binding site architecture, and evolutionary constraints. Generated sequences exhibited violations of natural genomic patterns and models showed particular difficulty with repetitive elements. To assess the quality of genome generation, we trained a convolutional neural network that reliably distinguished synthetic from natural sequences, achieving AUROC values up to 0.97 in eukaryotes and 0.82 in prokaryotes, with classification accuracy increasing monotonically with genomic distance from the seed. These findings suggest fundamental limitations in current gLM architectures for capturing the long-range, hierarchical nature of genomic sequences. Our work highlights the need for specialized architectures that explicitly model evolutionary constraints rather than relying solely on statistical patterns, with important implications for computational biology applications requiring realistic sequence generation and for biosafety assessments that depend on the distinguishability of synthetic and natural genomic sequences.

## Introduction

Large language models (LLMs) have revolutionized computational approaches across diverse domains, demonstrating unprecedented capabilities in understanding and generating complex sequential data ^1^. Their remarkable success in natural language processing has naturally prompted exploration of their potential for biological sequence analysis, particularly in computational biology where DNA, RNA, and protein sequences exhibit language-like properties with defined alphabets, syntactic rules, and hierarchical organization. The prospect of applying LLMs to genomic data is particularly compelling given the exponential growth of available sequence data and the increasing demand for computational tools capable of understanding complex genomic patterns, predicting functional elements, and generating biologically realistic sequences for synthetic biology applications ^2^.

Recent advances in genomic Language Models (gLMs) have achieved state-of-the-art performance across numerous biological tasks. As of early 2026, two main models possessing long-context genome generation capabilities are Evo 2^3^ and megaDNA^4^. Evo 2, released in February 2025, represents a significant milestone as the largest publicly available AI model for biology to date, featuring 40 billion parameters and trained on over 9.3 trillion nucleotides from approximately 128,000 genomes spanning all domains of life ^3^. This model can process sequences up to one million nucleotides at single-nucleotide resolution, enabling unprecedented long-range genomic modeling. MegaDNA is a more specialized 145-million-parameter model pre-trained on approximately 99,700 unannotated bacteriophage genomes with nucleotide-level tokenization, capable of generating de novo sequences up to 96 kilobases. These and other models^2^ have demonstrated strong performance in functional constraint and variant effect predictions, and sequence design tasks, achieving competitive or improved results in genome-wide variant effect prediction and regulatory element identification over computational methods specialized in these tasks. Concurrently, the rapid improvement of generative genomic models has raised questions about the potential biosafety implications of increasingly realistic synthetic genome generation, underscoring the need for rigorous model evaluations ^5^.

However, despite these encouraging developments, fundamental questions remain about the ability of current gLM architectures to faithfully capture the full complexity of genomic organization. Research on earlier gLMs revealed they primarily learned sequence patterns through recalling training data rather than understanding deeper contextual relationships ^6,7^. Unlike natural language, genomic sequences are sparse, highly repetitive, and often dominated by non-informative or low-information content regions, and are subject to intricate evolutionary constraints, functional dependencies, and multi-scale organizational principles that have been shaped by billions of years of selection. In this work, we systematically evaluate the limitations of state-of-the-art generative gLMs in reconstructing realistic genomic sequences. Specifically, we analyze Evo 2 ^3^ and megaDNA ^4,8^, which to our knowledge are the only current models designed for end-to-end genome-scale sequence generation. To enable rigorous assessment, we introduce a set of quantitative metrics that serve as benchmarks for comparing model-generated and natural genomic sequences. We reveal significant gaps between model-generated and natural genomic patterns that highlight the need for specialized architectures incorporating biological constraints. Furthermore, we show that a simple convolutional neural network is able to distinguish between natural and synthetically generated sequences, underscoring their discrepancies. Our findings suggest that while current approaches show promise for specific applications, substantial methodological advances are required before gLMs can reliably generate biologically realistic genomic sequences at scale.

## Results

The goal of this study was to systematically evaluate whether current genomic Language Models can generate biologically realistic genomic sequences that preserve essential organizational, compositional, and evolutionary constraints. When evaluating Evo 2, to capture the breadth of genomic diversity across the tree of life, we analyzed 200 complete genome assemblies from organisms spanning the major evolutionary lineages (**Methods; Supplementary Table 1**) and using several complementary metrics we compared synthetically generated genomes to their corresponding original genomes. In addition to the Evo 2 analyses, we evaluated megaDNA on bacteriophage genomes using two complementary comparison frameworks. For paired analyses, we analyzed 250 complete bacteriophage genomes, enabling direct comparison between natural sequences and megaDNA-generated counterparts. For population-level analyses, we used a large benchmark dataset comprising 4,969 natural bacteriophage genomes and 1,002 *de novo* synthetic bacteriophage genomes ^8^, allowing assessment of global compositional and structural trends in unpaired comparisons.

### Synthetic genomes fail to recapitulate organismal k-mer spectra

K-mer spectra describe the distribution of the frequencies of all possible k-length subsequences of a genome ^9^. In mammalian genomes, the k-mer spectrum often exhibits a bimodal (or more generally multimodal) distribution, plants show multimodal distributions, and most prokaryotes tend to show a unimodal spectrum ^9^. Because k-mer spectra also capture species-specific sequence organization, we wanted to examine the degree to which k-mer spectra of synthetic genomes capture these.

We analyzed representative species spanning major taxonomic groups (n=200), including tetrapods (*Homo sapiens*, *Mus musculus*), plants (*Oryza sativa*, *Zea mays*), and microbial and viral genomes. For each organism, we compared the k-mer spectra of the original genome to those of Evo 2-generated using the per-k-mer-type normalization of Chor *et al.* (2009) ^9^. We selected k following the heuristic *k* = ⌈0.7log_4_ *ℓ* ⌉; thus, pairwise 300 kbp windows used *k* = 7 and aggregated genome-scale analyses used *k* = 9; full details in Methods.

We observe that for all prokaryotic and eukaryotic species examined, the k-mer spectra of synthetic genomes are significantly different from the wild-type (one-sided Wilcoxon signed-rank test on window-level KS distances against Δ = 0.01, FDR-adjusted p-values < 0.0001) (**Supplementary Table 2**). Across all analyzed tetrapods, the wild-type genomes displayed the expected bimodal frequency distributions, while the Evo 2 synthetic genomes, exhibited a unimodal spectrum most of the time (**Fig 1A**, Kolmogorov-Smirnov, p-value<0.0001). In *Homo sapiens*, the synthetic sequences showed a systematic compression of the distribution: loss of rare k-mers (reduced diversity) and inflation of medium-frequency motifs (homogenization). The same pattern holds in *Mus musculus* and *Bos taurus*: wild-type spectra are bimodal, synthetic are unimodal and shifted toward intermediate abundances (**Fig 1C-F**, Kolmogorov-Smirnov, p-value<0.0001). For *Oryza sativa*, spectra appear closer visually, consistent with plants’ broader multimodality, yet Kolmogorov-Smirnov tests still detect significant discrepancies. Among prokaryotic and viral genomes, where wild-type spectra are naturally unimodal, Evo 2 outputs are qualitatively closer to the native distributions. Notably, both viral and prokaryotic genomes display an increased fraction of moderately rare k-mers, suggesting a systematic distortion of the k-mer spectrum (**Supplementary Figure 1**). We conclude that synthetic genomes generated by Evo 2 are currently unable to recapitulate the k-mer spectra of organismal genomes.

**Figure 1:**
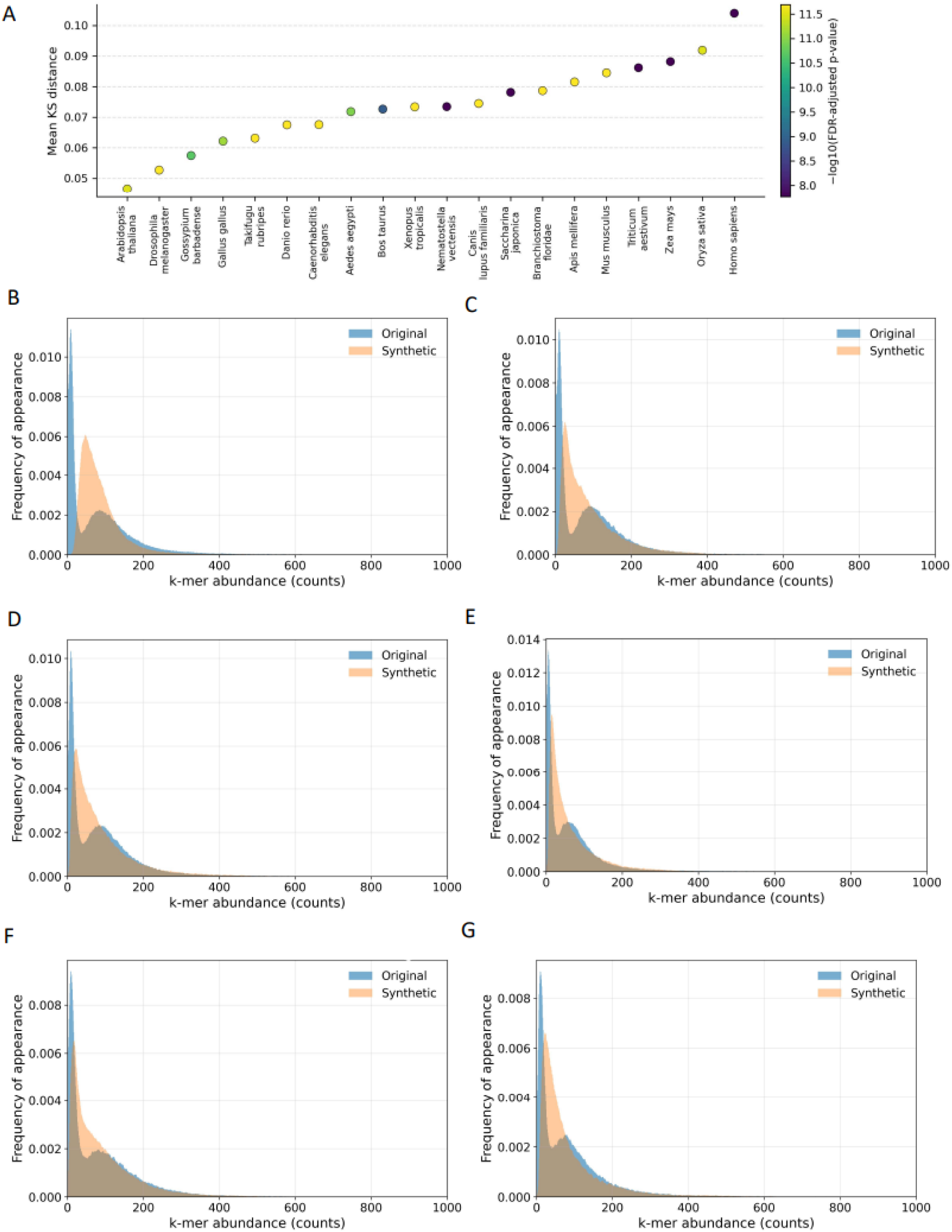
k-mer spectra of wild-type and synthetic genomes. **A.** K-mer spectrum bias levels between wild-type and synthetic genomes. **B-G.** Histograms of the k-mer spectra in **B** (*Homo sapiens*), **C** (*Mus musculus*), **D** (*Canis lupus familiaris*), **E** (*Bos taurus*), **F** (*Gallus gallus*), **G** (*Xenopus tropicalis*). k=7 for all plots.

### Chaos game representation of genomic k-mers in gLM-produced genomes

Chaos Game Representation (CGR) maps a genome sequence into a 2D plot by iteratively placing points based on successive k-mers, revealing patterns of nucleotide composition and structure ^10^. Each k-mer corresponds to a unique position in the plot, allowing CGR to visually and quantitatively compare genomes without sequence alignment. For each organismal genome studied, we generated CGR maps in original and synthetic genomes to study parity between them.

We observe that across the studied genomes, the synthetic counterparts displayed significantly different CGRs. This difference is quantitatively captured by frequency-based CGR (FCGR) analysis, which shows consistently elevated L1 distances between original and synthetic maps across taxa, accompanied by uniformly significant Wilcoxon signed-rank tests (all p < 5 × 10⁻⁴), indicating systematic shifts in higher-order k-mer spatial organization (**Fig 2A)**.

**Figure 2:**
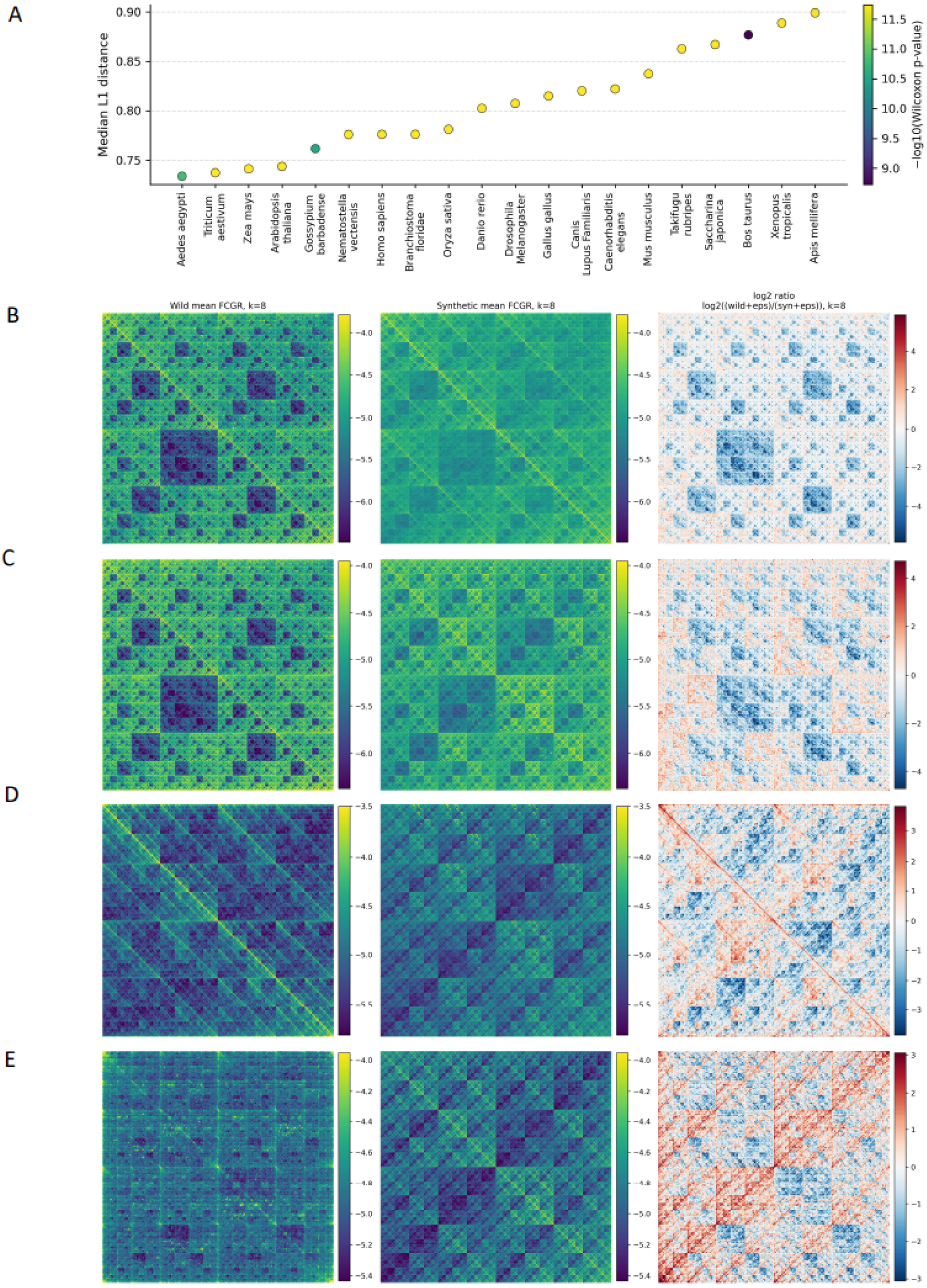
FCGR-based comparison of k-mer distributions between original and synthetic genomic sequences. **A.** Median L1 distances between normalized FCGRs of original and synthetic sequences across species, with color indicating Wilcoxon signed-rank significance (−log10 *p*). **B–E.** Mean FCGR tripanels (log10(P+ε)) for original (left) and synthetic (center) sequences, and their log2 ratio (right), shown for **(B)** *Homo sapiens*, **(C)** *Mus musculus*, **(D)** *Apis mellifera*, and **(E)** *Saccharina japonica*.

In all organisms examined, synthetic genomes exhibit systematic distortions in frequency chaos game representations (FCGRs) relative to their wild-type counterparts. FCGR distances are consistently large (median L1 distance 0.73-0.90 across eukaryotes), with highly significant Wilcoxon signed-rank tests across genome windows (all p < 3.2 × 10⁻⁸). Importantly, these differences do not reflect a uniform loss of spatial structure. Instead, synthetic genomes display a convergence toward a more homogenized FCGR pattern (**Fig 2B-E)**: in mammals, native multiscale contrast is dampened, whereas in taxa with inherently diffuse FCGRs, such as insects and algae, synthetic sequences exhibit artificially enhanced regularity. This indicates that Evo 2-generated genomes fail to preserve species-specific higher-order k-mer organization, instead converging toward an averaged k-mer frequency landscape.

### Failure of synthetic genomes to capture evolutionary nullomer constraints

Nullomers are short DNA sequences absent from a genome ^11,12^. Their absence from the genome has been linked to hypermutability and negative selection constraints ^13,14^. Here, we examined if there are differences in the nullomer content between wild-type and synthetic organismal genomes. We extracted the set of nullomers across all organismal genome assemblies (N=200) in our study. Due to difference in genome size, for viral genomes k-mer lengths between 4-10bp were examined, whereas in prokaryotes and eukaryotes the k-mer length thresholds were 7-10bp and 9-13bp respectively. We observe that synthetic genomes display a relative depletion in the number of nullomers identified for eukaryotes and bacteria.

Across 20 eukaryotic genomes, synthetic sequences showed consistent depletion in nullomer content relative to wild-type organismal genomes. We observed that the results were consistent across k-mer lengths (McNemar’s tests, multiple-testing correction using the Benjamini-Hochberg FDR), and the effect-size increased with increased k-mer length. Specifically, all species displayed statistically significant differences at k=10-13 (median q=0), whereas 45% of species displayed significant differences at k=9. Thus, we conclude that systematic differences in nullomer composition are essentially universal in eukaryotes once k ≥ 10 (**Fig 3A-I)**.

**Figure 3:**
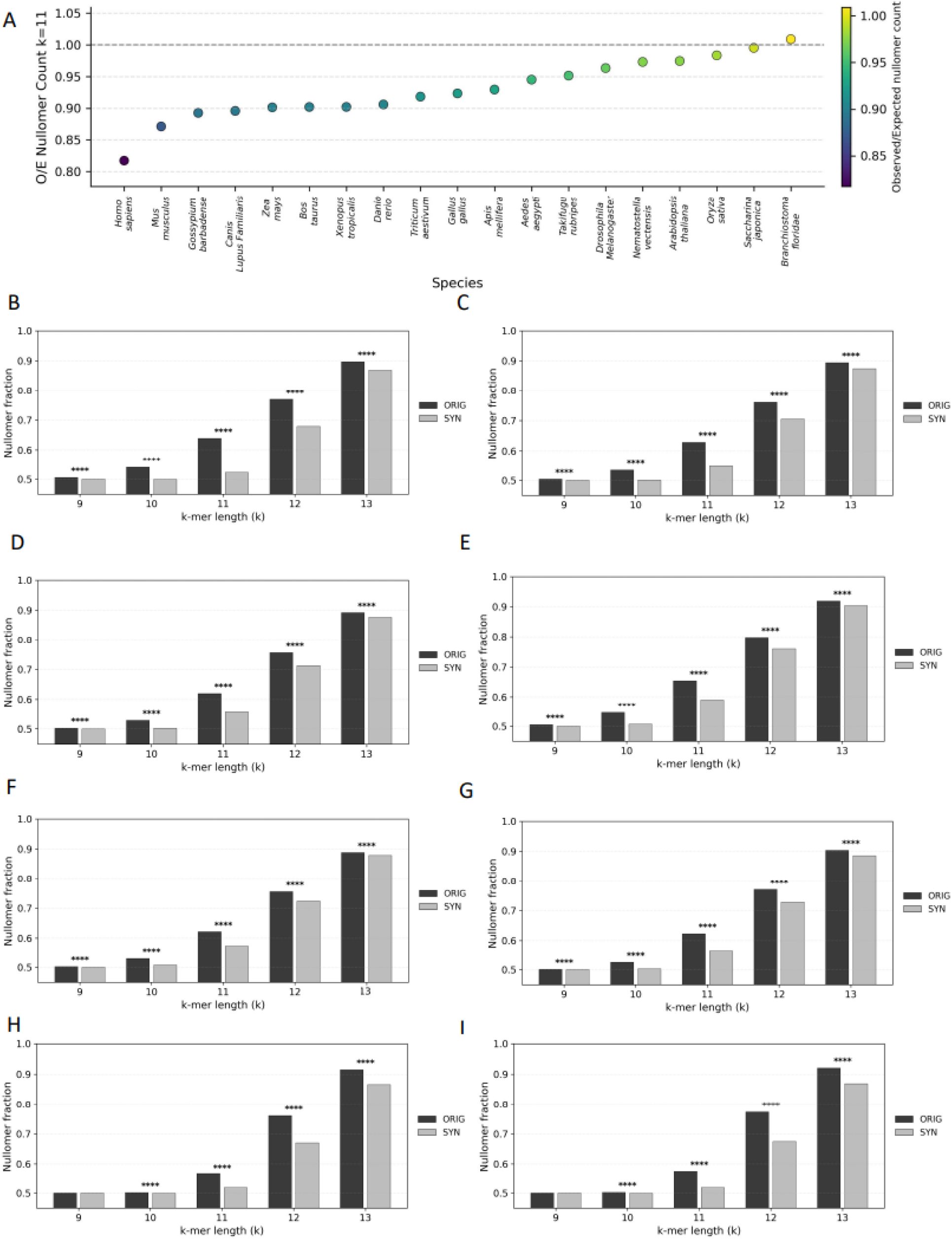
Relative depletion in nullomer content in synthetic genomes. **A-I. A** Number of observed (wild-type) and expected (synthetic) nullomers across organismal genomes. Histograms of nullomer content in wild-type and synthetic genomes in **B**. *Homo sapiens*, **C**. *Mus musculus*, **D.** *Canis lupus familiaris*, **E**. *Bos taurus*, **F**. *Gallus gallus*, **G**. *Xenopus tropicalis*, **H**. *Triticum aestivum*, and **I**. *Zea mays*. Adjusted p-values are displayed as * for p < 0.05, ** for p < 0.01, *** for p < 0.001, and **** for p < 0.0001.

In contrast to eukaryotic genomes, where synthetic sequences show a depletion of nullomers, viral and prokaryotic genomes exhibit an opposing pattern, underscoring the discrepancy between natural and synthetically generated nullomer patterns. Using k-mer presence-absence tests across the tested ranges (viruses: k = 4-10; prokaryotes: k = 8-12) with multiple-testing corrections, we observe significant, lineage-dependent divergence from wild type genomes. In both prokaryotes and viruses, synthetic genomes predominantly show an enrichment of nullomers relative to wild type, indicating altered k-mer exclusion patterns rather than preservation of native nullomer structure **(Fig S2A-D)**. The opposing trends observed between eukaryotic and viral or prokaryotic genomes likely reflect differences in genome architecture, specifically the sparsity of eukaryotic genomes, versus the compactness of viral and prokaryotic genomes, both of which the synthetic genomes cannot recapitulate. Together, these results suggest that Evo 2 imposes a largely domain-agnostic generative behavior that fails to capture organism-specific evolutionary constraints.

### Systematic distortion of non-B DNA motif landscapes in synthetic genomes

Non-B DNA structures are alternative conformations of the canonical right-handed B-form helix, including forms such as Z-DNA, G-quadruplexes, hairpins, cruciforms, H-DNA, and slipped or triplex motifs ^15^. These structures are biologically important across organismal genomes because they influence key genomic processes, such as replication, transcription, recombination, and repair, and are often hotspots for genomic instability ^15,16^. Regions that are predisposed to non-B DNA formation can be identified from the primary sequence. Given the prevalence of non-B DNA in organismal genomes and their biological roles, we examined if synthetic genomes were able to capture the same frequencies of non-B DNA sequences.

Across the 20 eukaryotic genomes analyzed, Evo 2-generated sequences exhibited a consistent depletion of non-B DNA motifs relative to their wild-type counterparts. By comparing matched regions and measuring how much more motif coverage was present in real genomes than in synthetic ones, we found that organismal genomes consistently showed higher motif coverage (**Fig 4A**). Direct repeats (DR), inverted repeats (IR), mirror repeats (MR), and short tandem repeats (STR) were depleted in 100% of eukaryotic genomes, while Z-DNA and G-quadruplexes (GQ) showed depletion in 95% and 85% of species, respectively. These depletions were frequently significant after multiple-testing correction (q < 0.05), with 90% of species being significant for DR and STR, 80% for IR, 80% for Z-DNA, 65% for MR, and 55% for GQ, indicating that the depletion trend is robust and reproducible across phylogenetically diverse eukaryotic genomes. Effect sizes were substantial and motif-class dependent. Median depletion factors (orig/syn) across species were 10.0-fold for DR, 5.63-fold for Z-DNA, 5.04-fold for STR, 2.64-fold for MR, 2.62-fold for GQ, and 1.77-fold for IR (**Fig. 4A**). Expressed inversely, synthetic sequences retained only ∼10% of wild-type DR coverage, ∼18% of Z-DNA, and ∼20% of STR at the median, underscoring a pronounced collapse of non-canonical structure-forming patterns in the generated sequence. While mean effects were inflated by outlier species (e.g., mean DR depletion 14.5-fold), the corresponding medians remained strongly shifted for every category, supporting a consistent cross-species reduction rather than a handful of outliers.

**Figure 4.**
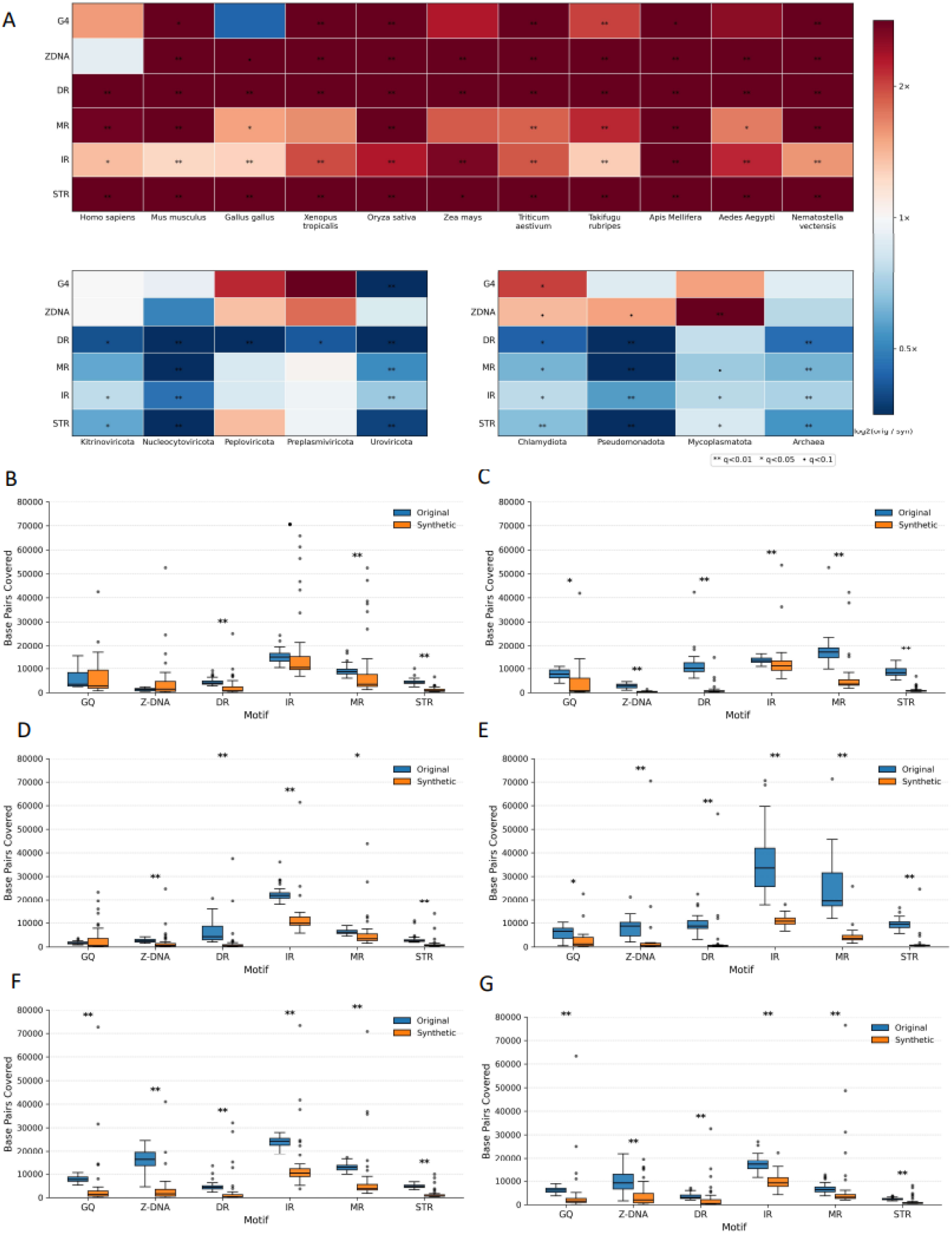
Comparison of non-B DNA motif content in original versus Evo-2-generated synthetic genomic sequences. **A.** Heatmap of median log₂(original/synthetic) base-pair coverage for six classes of non-B DNA motifs across representative eukaryotic species (top), viral groups (bottom left), and prokaryotic domains (bottom right). Positive values indicate depletion in Evo-2–generated synthetic genomes, whereas negative values indicate enrichment. Statistical significance after false discovery rate (FDR) correction is denoted by asterisks (* q < 0.1, ** q < 0.05, *** q < 0.01). **B-G.** Boxplots showing the distribution of base-pair coverage for six major non-B DNA motif classes, G-quadruplexes (GQ), Z-DNA, direct repeats (DR), inverted repeats (IR), mirror repeats (MR), and short tandem repeats (STR), in original (blue) and synthetic (orange) genomic windows. Panels correspond to the following species, in order: **B.** *Homo sapiens*, **C.** *Mus musculus*, **D.** *Aedes aegypti*, **E.** *Apis mellifera*, **F.** *Oryza sativa*, and **G.** *Triticum aestivum*. For each species, distributions are computed from matched 300-kbp genomic windows sampled proportionally to chromosome length. Values represent the total number of base pairs covered by each motif type per window. Statistical significance is indicated above each motif using FDR-corrected Wilcoxon signed-rank tests, with **q < 0.01** denoted by **, **q < 0.05** by *, and **q < 0.1** by •.

The most extreme depletions were observed in specific lineages or species and followed the same global direction (**Fig. 4B-G**). DR depletion was strongest in *Nematostella vectensis* (72.2-fold, q = 3.6×10⁻¹⁰) and *Saccharina japonica* (34.4-fold, q = 6.4×10⁻¹²), indicating near-elimination of DR-associated coverage in the synthetic sequences for these genomes. Similarly, the strongest GQ depletion occurred in *Saccharina japonica* (15.7-fold, q = 7.0×10⁻¹¹) and *Mus musculus* (9.28-fold, q = 2.1×10⁻²). Z-DNA showed the largest depletion in *Branchiostoma floridae* (12.6-fold, q = 1.4×10⁻⁸) and *Mus musculus* (12.6-fold, q = 2.5×10⁻³), while STR depletion peaked in *Saccharina japonica* (16.2-fold, q = 6.4×10⁻¹²) and *Apis mellifera* (14.1-fold, q = 3.2×10⁻¹¹) (**Fig. 4C**). For IR and MR, maximal depletions were smaller in magnitude but still substantial and significant (IR: *Apis mellifera* 2.87-fold, q = 1.3×10⁻¹¹; *Zea mays* 2.65-fold, q = 3.2×10⁻⁴; MR: *Apis mellifera* 5.64-fold, q = 4.1×10⁻⁸; *Mus musculus* 4.69-fold, q = 1.9×10⁻⁹) (**Fig. 4E**). Notably, significant enrichments in synthetic sequence were rare to absent at this resolution: the few species exhibiting nominal enrichment (negative log2 ratios) did not pass FDR, reinforcing that the dominant and statistically supported pattern in eukaryotes is depletion rather than motif inflation.

In archaea, non-B DNA motifs showed a strong and consistent enrichment in synthetic genomes, with significant effects detected in ∼83% of motif tests (**Fig. 4A**). All repeat-based motifs (DR, IR, MR, STR) as well as Z-DNA were significantly enriched in synthetic sequences, whereas GQs showed no significant difference despite a similar enrichment trend. In bacteria, non-B DNA motif differences showed strong motif-specific patterns, with significant effects in ∼72% of tests (**Fig. 4A**). Repeat-based motifs (DR, IR, MR, STR) were consistently and significantly enriched in synthetic genomes across all bacterial phyla, whereas GQs and Z-DNA displayed mixed, often depleted, and less consistent signals. In viral genomes, non-B DNA motifs showed heterogeneous but motif-specific differences between synthetic and wild-type sequences. Significant effects were detected in ∼47% of all virus-motif tests, driven primarily by DRs, IRs, MRs, and STRs, whereas GQs and Z-DNA showed little to no consistent signal. When significant, effects were predominantly enrichments in synthetic genomes, with DR exhibiting the strongest and most consistent signal (80% significant across viral groups), followed by IR, MR, and STR (each 60%) (**Fig. 4A**).

Together, these results demonstrate that Evo 2-generated eukaryotic sequences systematically under-represent non-B DNA-forming motifs, with the strongest losses affecting repeat-associated categories (DR and STR) and substantial, widespread depletion also extending to Z-DNA and GQ. This consistent directionality across animals, plants, and diverse eukaryotic lineages suggests that the generative process preferentially produces sequence compositions that reduce the occurrence of repetitive and structure-prone substrings, potentially reflecting implicit constraints learned from training data, generation-time biases, or a tendency toward “regularized” sequence outputs that avoid motif-dense regions. Unlike eukaryotes where non-B DNA motifs are broadly depleted, viral synthetic genomes show an excess of non-B DNA categories. Overall, synthetic genome generation by Evo 2 consistently fails to replicate the non-B DNA density in organismal genomes, particularly in large and repeat-rich genomes.

### Transcription factor binding sites are systematically enriched in synthetic genomes

To assess whether Evo 2-generated human sequences preserve regulatory motif composition, we analyzed 1,019 transcription factor profiles and compared transcription factor binding site (TFBS) repertoires between wild-type sequences and their synthetic counterparts.

TFBS analysis revealed pronounced and systematic distortions, with the vast majority of transcription factors showing higher TFBS abundance in synthetic sequences relative to the original human genome (**Fig. 5A**, binomial test, p-value<0.0001). This asymmetry suggests a systematic enrichment of transcription factor binding motifs in Evo2-generated sequences rather than random fluctuation around parity. Several motifs exhibit extreme effect sizes and statistical significance, implying that Evo 2 does not merely introduce noise but actively reshapes the regulatory motif landscape, increasing the density of specific TFBS beyond levels observed in native human genomic regions. Importantly, this enrichment is motif-dependent, with some transcription factors being strongly overrepresented while others remain closer to parity, indicating selective distortion rather than uniform inflation.

**Figure 5:**
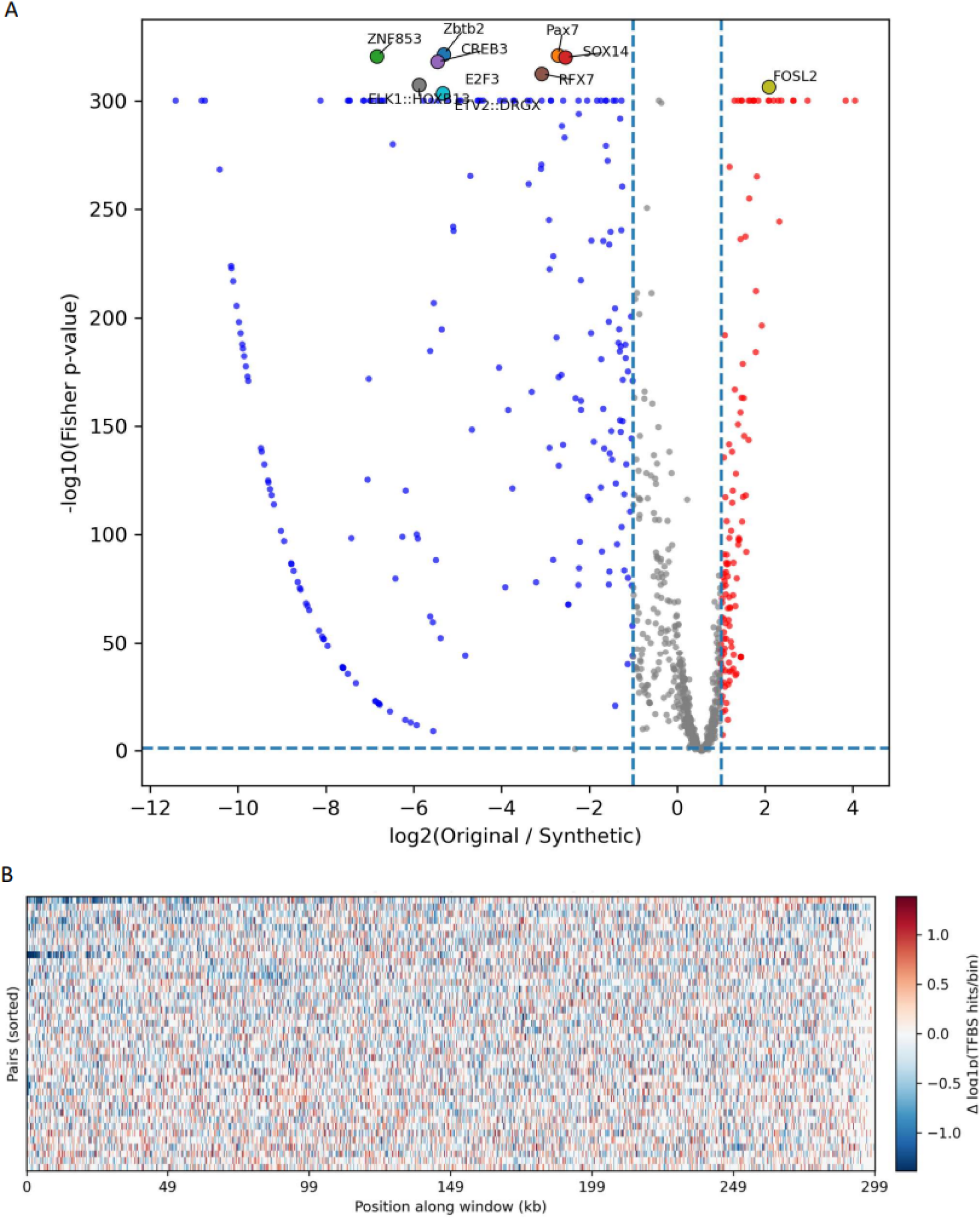
TFBS enrichment and spatial redistribution in synthetic human genomes. **A.** Volcano plot of motif-level TFBS differences between original and synthetic human genomic windows. The x-axis shows log2(original/synthetic) TFBS counts per motif and the y-axis shows −log10(Fisher’s exact test p-value). **B.** Heatmap of positional TFBS density differences (log1p-scaled; original − synthetic) along 300 kb windows, computed using the ten most significant motifs from panel A.

Beyond overall motif abundance, we examined whether Evo 2 preserves the spatial organization of TFBS along human genomic windows. We find that in wild-type sequences, TFBS tend to concentrate into localized high-density regions, whereas synthetic sequences display a comparatively more uniform distribution along the window length (**Fig. 5B**). This effect was quantified using complementary clustering metrics computed per window pair: both the variance-to-mean ratio (Fano factor) and the Gini coefficient were significantly higher in original sequences than in synthetic ones (paired Wilcoxon p ≤ 10⁻⁶), indicating a loss of TFBS clustering in synthetic genomes. By contrast, short-range spatial autocorrelation which captures whether nearby bins tend to share similar TFBS density, did not differ significantly between original and synthetic sequences. This suggests that Evo 2 does not randomize local TFBS placement, but instead reduces the intensity of large TFBS hotspots, leading to a more even distribution along the window. We conclude that Evo 2–generated human sequences exhibit systematic, motif-dependent enrichment of transcription factor binding sites and a pronounced loss of native clustering and hotspot organization.

### Compositional and structural divergence in megaDNA-generated phage genomes

To assess whether the failures observed across our benchmarks for Evo 2 generalize to other genome-scale generative models, we evaluated megaDNA on bacteriophage sequences. Unlike Evo 2, megaDNA is trained on a narrower domain and operates at shorter generation lengths (96 kb), providing a complementary test case for sequence realism under more constrained settings. We evaluated megaDNA on bacteriophage genomes using a paired analysis of 250 complete genomes following a similar methodology to our Evo 2 analyses. We also used the dataset of 1,002 *de novo* synthetic and 4,969 natural bacteriophage genomes from ref ^8^, which previously showed that natural and megaDNA-generated sequences are compositionally distinct on a separate set of metrics. We report that using population-level analyses on this dataset (see methods) our results on our set of metrics conform with the previously reported differences reported by ref ^8^.

MegaDNA-generated phage genomes exhibited distortions in k-mer spectra relative to their natural counterparts. Across paired comparisons, synthetic sequences preserved the overall shape of the native spectrum but showed a systematic redistribution of k-mer mass toward moderately rare and intermediate-abundance k-mers (median KS ≈ 0.072; paired tests p = 4.62×10⁻⁴³, FDR-matched; **Fig. 6A-B**). Population-level analyses revealed a strong length dependence of this effect. Specifically, the shortest phage genomes showed the most pronounced collapse of k-mer support and the intermediate and larger genomes displayed subtler internal reshaping with little net gain or loss of total k-mer mass (**Fig. 7A**). We also evaluated higher-order compositional organization using FCGR. Despite partial preservation of k-mer frequency spectra, FCGRs showed a consistent divergence between natural and megaDNA-generated phage genomes across all paired comparisons and genome length bins, indicating a pervasive disruption of k-mer organization independent of genome length. In paired analyses, the separation was substantial (median L1 distance ≈ 1.10) and statistically significant under paired testing (p = 4.63 × 10⁻⁴³; **Fig. 6C**). Consistently, population-level comparisons across genome length bins showed pronounced FCGR divergence (bin-wise L1 range ≈ 0.40–0.93), with all bins remaining significant under permutation testing after multiple-testing correction (q_FDR ≤ 0.0014; **Fig. 7C**).

**Figure 6:**
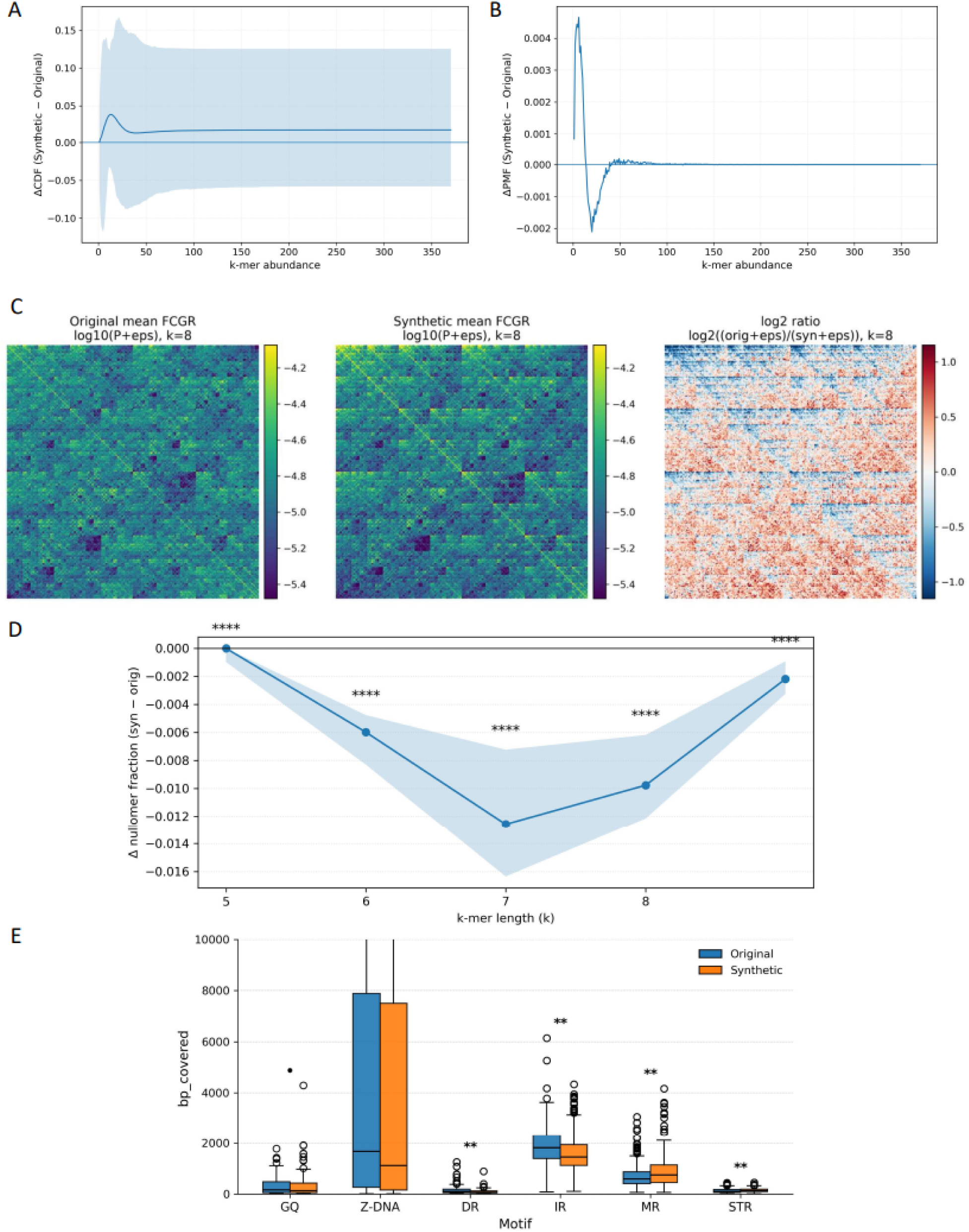
Compositional and structural deviations in megaDNA-generated bacteriophage sequences. **A.** Difference in k-mer abundance cumulative distribution functions (ΔCDF = synthetic - original) computed across paired wild-type and megaDNA-generated phage windows, showing systematic distortions in k-mer frequency distributions. Shaded regions indicate variability across pairs. **B.** Difference in the mean k-mer abundance probability mass function (ΔPMF = synthetic − original), highlighting abundance-specific redistribution of distinct k-mers between natural and synthetic sequences. **C.** Frequency chaos game representations (FCGR, k = 8) comparing mean original and synthetic k-mer frequency landscapes (log₁₀-scaled), alongside the log₂ ratio map highlighting structured frequency shifts between wild-type and generated sequences. **D.** Median change in nullomer fraction (Δ = synthetic − original) across k-mer lengths (k = 5-8), with 95% bootstrap confidence intervals. Significance denotes deviation from zero based on a Wilcoxon signed-rank test across paired windows, with false discovery rate correction applied across k. **E.** Coverage of predicted non-B DNA motifs (bp covered) in original and synthetic phage sequences, shown as paired distributions across motif classes. Asterisks denote statistically significant differences between original and synthetic sequences (*p < 0.05, **p < 0.01, ****p < 1e-4).

**Figure 7:**
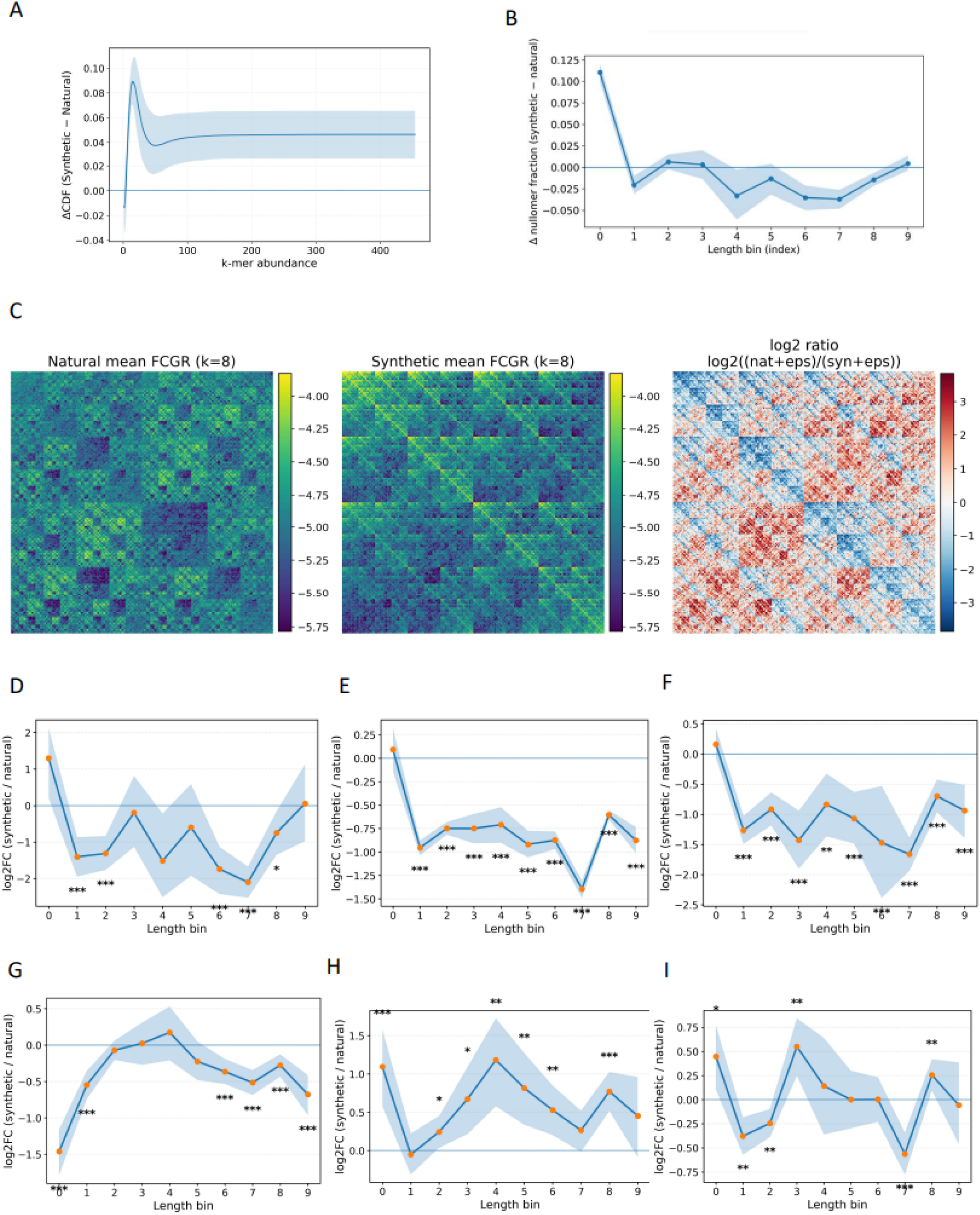
Length-stratified differences between natural and synthetic phage genomes across multiple sequence descriptors. **A.** Difference in cumulative distribution functions (ΔCDF = synthetic − natural) of k-mer spectra for k = 6 in the 50–57 kb length bin, shown as a function of k-mer abundance. Shaded region denotes the 95% bootstrap confidence interval. **B.** Difference in nullomer fraction between synthetic and natural genomes across length bins, with positive values indicating enrichment in synthetic sequences. Points show mean differences and shaded regions indicate 95% bootstrap confidence intervals. **C.** Frequency chaos game representations (FCGR; k = 8) for genomes in the 50–57 kb bin: natural mean FCGR (left), synthetic mean FCGR (center), and log2 ratio map (log2[(natural + ε)/(synthetic + ε)]) highlighting systematic spatial differences in higher-order k-mer organization. **D–I.** Log2 fold-change in non-B DNA motif coverage between synthetic and natural genomes (log2FC = log2[syn/nat]) across length bins, computed on balanced per-bin samples. Shaded regions indicate 95% bootstrap confidence intervals. Panels show results for **(D)** G-quadruplexes (G4Hunter), **(E)** Z-DNA (ZSeeker), **(F)** direct repeats (DR), **(G)** inverted repeats (IR), **(H)** mirror repeats (MR), and **(I)** short tandem repeats (STR). Asterisks denote bins with significant differences after permutation testing and FDR correction (q < 0.05).

Moreover, we examined whether megaDNA preserves native k-mer exclusion patterns using nullomer analysis. In paired comparisons, synthetic phage genomes consistently showed reduced nullomer content relative to their natural counterparts, indicating systematic filling-in of k-mers that are absent from wild-type sequences. This depletion was strongest at intermediate k values (around k = 7) and was statistically significant under FDR-corrected paired testing (**Fig. 6D**). Population-level analyses further revealed that this effect depends on both genome length and kmer length, with distinct regimes of nullomer depletion and enrichment across length classes rather than a single consistent scaling behavior (**Fig. 7B**). Finally, we assessed whether megaDNA preserves native non-B DNA motif patterns. Across motif classes, synthetic phage genomes showed systematic and motif-specific deviations in non-B DNA coverage relative to natural sequences in both paired and population-level analyses (**Fig. 6E**, **Fig. 7D-I**). In paired comparisons, direct and inverted repeats were consistently depleted in synthetic genomes, with large positive log₂ fold changes (natural > synthetic) and strong statistical support (DR: q ≈ 1.7 × 10⁻¹²; IR: q ≈ 1.6 × 10⁻⁹), indicating loss of native repeat-associated non-B structure. By contrast, mirror repeats and short tandem repeats were significantly enriched in synthetic genomes (MR: q ≈ 6.7 × 10⁻⁶; STR: q ≈ 1.0 × 10⁻⁵). Z-DNA and G-quadruplex motifs showed weaker effects in paired analyses, with Z-DNA largely unchanged and G4 displaying only borderline shifts.

Population-level analyses across genome length bins reinforced these patterns while revealing length-dependent modulation. Direct repeats were consistently depleted across nearly all bins (synthetic retaining ∼31-62% of natural coverage; q_FDR ≤ 0.005), whereas mirror repeats were broadly enriched, reaching ∼1.4-2.3-fold higher coverage in synthetic genomes across multiple bins (q_FDR ≤ 0.043). Z-DNA showed the most consistent population-level depletion signal, with synthetic genomes retaining ∼38–66% of natural coverage across bins (median ≈ 55%; q_FDR ≈ 5.6 × 10⁻⁵), while G-quadruplexes were also depleted in most intermediate bins (retaining ∼23–60% of natural coverage; q_FDR ≤ 0.025). Inverted repeats and STRs exhibited mixed, length-dependent behavior, with significant effects confined to specific length regimes.

Together, our findings show that megaDNA produces phage genomes that disrupt higher-order compositional organization, nullomer content, and non-B DNA motif landscapes in a length- and motif-dependent manner.

### CNN model reliably distinguishes natural and synthetically generated genomes

We trained a convolutional neural network model to distinguish synthetic from natural sequences based on chunks of 1024 bp. In eukaryotes, AUROC increased from 0.89 with a single chunk to 0.97-0.98 with ≥8 chunks, while F1 rose from 0.80 to ∼0.93. Performance in prokaryotes was lower overall but showed a similar aggregation trend, with AUROC increasing from ∼0.74 (1 chunk) to ∼0.82 (16 chunks), and F1 peaking at ∼0.74 before declining slightly at 32 chunks. Viral classification exhibited more modest discrimination, with AUROC ranging from ∼0.62 (1 chunk) to ∼0.69 (16 chunks) and F1 from ∼0.55 to ∼0.61. Overall, these results demonstrate that even a relatively simple CNN, without handcrafted genomic features, can robustly distinguish synthetic from natural DNA (**Fig. 8A–B**). Notably, discrimination is strongest in eukaryotes and weaker in viruses, suggesting that synthetic sequence generation more readily diverges from the structure of complex genomes. In contrast, lower-complexity genomes appear more difficult to distinguish, indicating that current generative models more closely approximate their sequence-level characteristics.

**Figure 8.**
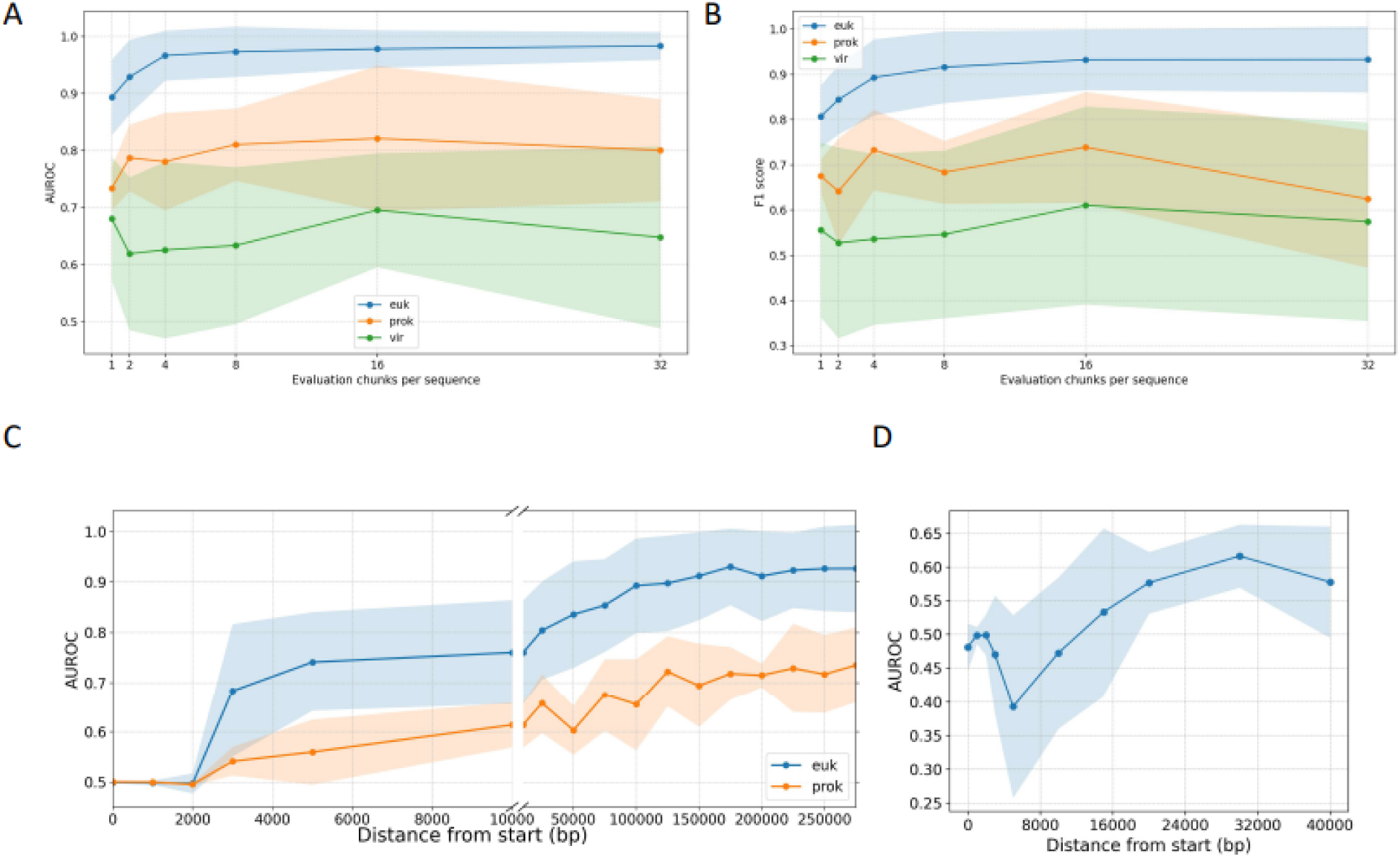
Classification performance for distinguishing synthetic from natural sequences across taxonomic domains. (**A–B**) AUROC (**A**) and F1 score (**B**) as a function of the number of evaluation chunks per sequence for eukaryotes (euk), prokaryotes (prok), and viruses (vir), using a CNN trained to classify sequences as synthetic or natural. Solid lines indicate mean performance across leave-one-tag-out evaluations; shaded regions represent ±1 standard deviation across holdout tags. (**C–D**) AUROC as a function of genomic distance from the seed region used for sequence generation (single-chunk evaluation), shown for eukaryotic and prokaryotic domains (**C**) and for the viral domain (**D**). Shaded regions denote ±1 standard deviation across tags.

Το further understand the discrepancies between natural and generated genomes, we evaluated our classifier’s performance across evaluation chunks that were found in different distances from the original seed (prompt) of the gLM. Classifier performance increased monotonically with distance from the 3 kb seed (**Fig. 8C–D**), revealing clear degradation with increased context length, matching results previously reported in Large Language Models ^17^. In eukaryotes, AUROC was indistinguishable from chance near the seed (0–2 kb; AUROC ≈ 0.50), but rose sharply beyond 3 kb (AUROC 0.68 at 3 kb, 0.76 at 10 kb), reaching >0.90 beyond 150 kb and peaking at 0.93 at 175 kb. A similar but attenuated pattern was observed in prokaryotes. AUROC was ∼0.50 at 0-2 kb, increased gradually with distance (0.61 at 10 kb, 0.66 at 25 kb), and reached ∼0.73 at 275 kb. Viruses exhibited the same qualitative trend over a much shorter genomic scale: AUROC was near chance close to the seed (∼0.48–0.50 within the first 2 kb), increased progressively with distance, and reached ∼0.62 by 30 kb before slightly declining at the most distal position. The fact that discrimination is near random within close proximity to the conditioning window but becomes progressively stronger at distal positions indicates that synthetic sequences preserve local statistics but increasingly diverge from natural genomic structure over long genomic distances. This progressive degradation provides quantitative evidence of long-range context collapse in current generative genome models.

## Methods

### Genome selection

For Evo 2, we generated 200 synthetic organismal genomes, spanning different taxonomic groups, including vertebrates, invertebrates, plants, fungi, algae, and protozoa. Complete genomes used as controls were downloaded from the GenBank and RefSeq databases ^18,19^. These included N=20 eukaryotic, N=52 bacterial, N=9 archaeal and N=129 viral organismal genome assemblies (**Supplementary Table 1)**. The selection covered both repeat-rich genomes such as vertebrates and flowering plants, as well as compact genomes such as yeast, nematodes, and apicomplexans. We selected representative phyla for viral (Nucleocytoviricota, Peploviricota, Uroviricota) and prokaryotic groups (Archaea, Chlamydiota, Mycoplasmatota, Pseudomonadota).

MegaDNA was evaluated using two complementary comparison frameworks: paired, genome-matched analyses and unpaired, population-level comparisons. For population-level analyses, we used the publicly available bacteriophage benchmark dataset from Transformer model generated bacteriophage genomes are compositionally distinct from natural sequences ^8^, which comprises 4,969 natural bacteriophage genomes and 1,002 de novo synthetic bacteriophage genomes.

For paired analyses, we assembled a non-redundant set of 250 natural bacteriophage genomes to serve as reference sequences for megaDNA generation. This set included (i) 49 bacteriophage genomes drawn from the viral genomes analyzed in the Evo 2 evaluation, (ii) 50 additional bacteriophage genomes randomly sampled from NCBI (Supplementary Table 1), and (iii) 151 bacteriophage genomes drawn from the same benchmark dataset described above^8^. Together, these 250 genomes span a broad range of genome lengths and were used for paired comparisons between natural and megaDNA-generated sequences.

### Synthetic genomes generation

For Evo 2, from each eukaryotic reference genome assembly, we sampled N = 40 windows of 300,000 bp proportionally to chromosome length. This window size was chosen based on a preliminary analysis of long-range generation stability, which revealed low-complexity collapse during extended Evo 2 generations, characterized by multi-kilobase homopolymer runs (e.g., ≥5–10 kb of a single nucleotide). Such events occurred after ∼300–400 kb of autoregressive decoding, motivating the use of fixed-length 300 kb windows to limit collapse while preserving long-range context. Sampling was restricted to primary chromosomes (NC_* or canonical chr names), enforced a 10,000 bp minimum inter-window gap on the same chromosome and used an N-content threshold of 10%. Sampling was deterministic with seed = 1337. Accepted windows were saved as FASTA records named by genomic coordinates. For each 300 kbp window, Evo 2 was prompted with a species-specific phylogenetic tag (phylotag) followed by the first 3,000 bp of the native window (“seed”), and asked to generate additional bases to reach exactly 300,000 bp. Decoding hyperparameters were fixed across all runs: temperature = 1 and top-k = 4. Model outputs were filtered to the DNA alphabet (A/C/G/T/N). For each window we retained the native FASTA, the Evo 2 synthetic FASTA, and a manifest row linking the pair for downstream analyses. For genomes with total length ≤ 300 kbp, we did not sample windows. Instead, we used the same prompting and decoding scheme as above, phylotag + first 3,000 bp seed, temperature = 1, top-k = 4 and generated additional bases to reach the full native genome length in a single shot. Outputs were filtered/padded as above to match the exact target length.

For megaDNA, synthetic bacteriophage sequences were generated using the pretrained 145M-parameter megaDNA model under a seed-conditioned autoregressive decoding scheme. For each natural reference genome used in the paired analyses, the first 3,000 bp were provided as a fixed conditioning seed, and the model was instructed to generate additional sequence to a target length of 50,000 bp. This configuration was chosen to balance sufficient sequence length for detecting higher-order compositional and structural deviations while avoiding excessive truncation or collapse effects in long-range generation. Sequence generation was performed with a sampling temperature of 0.95 and no token probability filtering (filter threshold = 0.0), following the inference settings reported in the original megaDNA study.

### K-mer spectra analysis

To compare the compositional properties of the original and synthetic sequences, we computed k-mer frequency spectra following the approach of Chor et al. (2009) ^9^. For each pair of original and synthetic sequences, all possible k-mers were counted separately per complete genome assembly to ensure contig-aware processing (i.e., no k-mers crossed record boundaries). Only canonical DNA bases (A, C, G, T) were considered, and lowercase bases were treated as masked positions.

The value of k was determined automatically from the effective sequence length using the heuristic *k* = 0. 7 × *log*_4_ (*lengt*ℎ), where length denotes the number of valid A/C/G/T bases in the shorter of the two sequences. For each sequence, we built a per-k-mer-type probability distribution by normalizing the abundance histogram over the total k-mer space (4^k^). We visualized these distributions as i) spectra plots, showing the normalized frequency of distinct k-mers at each abundance level, ii) cumulative distribution functions (CDFs), displaying cumulative k-mer fractions, and iii) Quantile–quantile (Q–Q) plots, comparing k-mer abundance quantiles between original and synthetic genomes. To quantify similarity between spectra, we computed three non-parametric metrics: i) Kolmogorov-Smirnov (KS) statistic and p-value to test equality of the two cumulative distributions, ii) Jensen-Shannon divergence (JSD) to measure information-theoretic distance, and iii) Earth Mover’s Distance (EMD) as the L1 difference between cumulative distributions. All results were aggregated across species to assess global compositional similarity between real and generated genomic sequences, providing quantitative metrics for synthetic sequence detectability..

### Comparison of nullomer landscapes between natural and Evo 2-simulated genomes

To quantify the depletion of specific sequence patterns in natural versus gLM-generated genomes, we analyzed nullomers. Nullomer detection was performed using the KMC k-mer counting suite (v3) ^20^, applied to concatenated FASTA files representing all original and synthetic windows per organism. KMC was executed for k-mer lengths k=4-10bp for viral genomes, 8-12bp for prokaryotes, and 9-13bp for eukaryotes. For each k, we extracted the number of distinct observed k-mers from the KMC histograms. The total possible k-mer space (4^k) was used to compute the number of missing k-mers (nullomers) and their relative fraction:

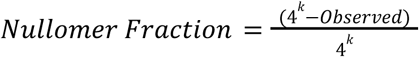

Boxplots and barplots were generated to visualize nullomer counts and fractions across k values (11-15 bps). Δ-fraction plots were also produced, showing the change in nullomer fraction between synthetic and original genomes. To assess statistical significance, we aggregated results across all analyzed species and performed Wilcoxon signed-rank tests (paired across species) comparing nullomer fractions between original and synthetic genomes for each k. Significance levels were reported per k, and trends in mean Δ values were interpreted as indicative of systematic overrepresentation or depletion of specific k-mers in gLM 2-generated genomes relative to their natural counterparts.

### Non-B DNA motif analysis

Non-B DNA structural motifs were analyzed in both the original and synthetic sequences using three specialized tools: Zseeker ^21^, G4Hunter ^22^, and non-B GFA ^23^. Each pair of sequences was processed independently, and results were harmonized to quantify motif counts and total base-pair coverage per motif type. Potential G-quadruplex-forming sequences were identified using G4Hunter, run with default parameters (window size = 25 bp, score threshold = ±1.5). Both the number of G-quadruplex motifs and their cumulative coverage were recorded. Potential Z-DNA forming sequences were detected with ZSeeker, using default parameters. Additional non-B sequences, including direct repeats, inverted repeats, mirror repeats, and short tandem repeats, were detected using the non-B DNA GFA tool (C binary) with default parameters.

For each non-B DNA motif type, we computed two quantitative metrics per sequence: (i) the number of detected loci (n_hits) and (ii) the total number of base pairs covered (bp_covered). Results were summarized as boxplots comparing the distributions between original and synthetic genomes. Statistical significance of differences was evaluated using paired t-tests and Wilcoxon signed-rank tests with multiple testing correction. Finally, to capture large-scale compositional trends across species and domains, we calculated the log₂ ratio of base-pair coverage (orig/syn) for each motif and aggregated the median log₂(orig/syn) values across all genomes. These values were visualized as three-panel heatmaps displaying median motif enrichment or depletion for eukaryotic, viral, and bacterial phyla. Each cell in the heatmap represents the median log₂(orig/syn) of base-pair coverage for a specific motif and taxonomic group. Significance was assessed per cell using Wilcoxon signed-rank tests against zero, followed by Benjamini-Hochberg FDR correction within each panel. Significant cells were annotated with q < 0.1 (*), q < 0.05 (*), or q < 0.01 (**).

### TFBS scanning in human genomic windows

TFBSs were identified exclusively in human genomic windows using FIMO from the MEME suite ^24^ and transcription factor motifs from the JASPAR 2026 vertebrates database ^25^. For each original-synthetic window pair, both sequences were scanned using identical parameters. Motif occurrences were filtered using a site-level threshold of p-value < 1e^−4^, yielding high-confidence TFBS calls for downstream analysis.

### Motif-level aggregation and statistical testing

Filtered TFBS hits were aggregated across all human window pairs on a per-motif basis. For each motif *m*, we computed the total number of hits in original human sequences (H_orig,m) and in synthetic sequences (H_syn,m), along with the total number of hits across all motifs in each condition. Motif-specific enrichment was evaluated using Fisher’s exact test applied to a 2×2 contingency table contrasting motif-specific and background hit counts between original and synthetic sequences.

Effect sizes were summarized using the log_2_ fold-change:

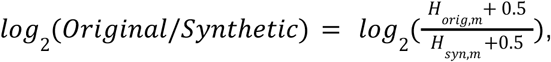

where a pseudocount of 0.5 was added to avoid undefined ratios. Negative values therefore indicate motif enrichment in synthetic sequences, while positive values indicate enrichment in the original human genome.

Motif-level results were visualized using a volcano plot, with −log_10_(Fisher p-value) plotted against log2(original/synthetic). For visualization purposes, motifs with p < 0.05 and |log2FC| > 1 were highlighted, and the top motifs by statistical significance were emphasized in the legend (Fig. 5A). This representation highlights the strong skew toward synthetic-enriched TFBSs in human Evo2-generated sequences.

For positional analysis, TFBS hits were binned along each 300 kbp human window and converted to log1p-scaled densities for original and synthetic sequences. Per-bin differences were computed as:

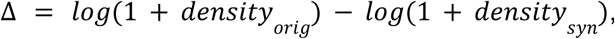

and visualized as a heatmap across all window pairs. Negative values indicate higher TFBS density in synthetic sequences. This analysis enables assessment of whether TFBS enrichment is localized or distributed across window positions.

### MegaDNA population-level tests

Natural and synthetic phage genomes were not paired and spanned a wide range of lengths. To enable meaningful comparison, genomes were first stratified by length into discrete bins, and all analyses were conducted within bins to control for length-dependent effects. Comparisons were therefore made between distributions of natural and synthetic genomes matched by length class rather than by one-to-one pairing.

Because the number of genomes differed between natural and synthetic sets within each bin, we applied within-bin balancing by randomly downsampling the larger group to match the smaller one. All reported effect sizes and statistical tests were computed on these balanced samples, ensuring that results were not driven by unequal sample sizes. This balancing procedure was applied independently for each bin and repeated implicitly within resampling procedures.

Effect sizes were computed at the per-genome level and then aggregated within bins. Uncertainty was estimated using nonparametric bootstrap resampling of genomes within each group to derive confidence intervals. Statistical significance was assessed using two-sided permutation tests performed within each bin by randomly shuffling natural/synthetic labels across pooled genomes. To account for multiple testing across bins (and motif classes where applicable), p-values were adjusted using the Benjamini–Hochberg false discovery rate (FDR) procedure. All analyses were performed on identically stratified and balanced data, allowing direct comparison of effect magnitudes and significance patterns across the different sequence-level descriptors.

### Wild versus synthetic sequence classification

To directly assess whether synthetic genomes are computationally distinguishable from natural sequences, we trained supervised dilated convolutional neural network (CNN) classifiers to predict whether a genomic window originated from a wild-type or model-generated genome. For each taxonomic domain (eukaryotes, prokaryotes, and viruses), balanced datasets of fixed-length windows were constructed from natural genomes and their corresponding synthetic counterparts. To assess cross-taxon generalization, we employed a leave-one-group-out scheme. For eukaryotes, each run held out one species during training and evaluated on the excluded species; this was repeated across all species, and performance was reported as mean ± standard deviation. For prokaryotes and viruses, the same procedure was applied at the phylum level. Sequences were one-hot encoded (A/C/G/T) and provided as input to the model. During training, each sequence was represented by N randomly sampled chunks of length 1024bp, and chunk-level logits were averaged to obtain a sequence-level prediction. At evaluation, the number of sampled chunks per sequence was varied (1, 2, 4, 8, 16, or 32), and logits were averaged to produce the final sequence-level score. The architecture consisted of stacked 1D dilated convolutional layers with increasing dilation rates, enabling the network to capture both local motif patterns and long-range dependencies, followed by fully connected layers and a sigmoid output for binary classification.

To quantify how discriminability varies with distance from the conditioning seed, we performed a distance-from-seed evaluation on the held-out test tag. After training with random chunk sampling, we scored each test sequence using fixed-position chunks of length 1024 starting at predefined offsets from the sequence start (e.g., 0, 1, 2, 3, 5, 10 kb), and computed AUROC/F1 as a function of offset, aggregating results across holdout folds.

Models were trained separately for eukaryotic, prokaryotic, and viral datasets to account for domain-specific genome architecture. Specifically, for viral classification, synthetic sequences included both Evo 2–generated viral genomes and megaDNA-generated bacteriophage genomes. We optimized hyperparameters through systematic search on validation sets. Model performance was evaluated on held-out test sets using AUROC and F-1 score.

## Discussion

Our study demonstrates that current state-of-the-art large language model architectures fail to recapitulate essential principles of organismal genomes. We examined the ability of Evo 2 and megaDNA to recapitulate basic characteristics of organismal genomes including their k-mer spectra, their nullomer profiles, their chaos game representations, their non-B DNA content and the TFBS content. gLM-generated genomes displayed pervasive distortions across these properties, including biased k-mer spectra, loss of long-range compositional heterogeneity, and failure to reproduce an accurate regulatory TFBS profile in the human genome. More broadly, our results suggest that Evo 2 and megaDNA trained on genomic sequences struggle to internalize the fundamental principles of genome organization. These include the sparsity of functional signal, the high noise-to-signal ratio, the redundancy in sequence composition, and the diverse evolutionary constraints that shape genomes, properties that differ profoundly from the structured and semantically dense patterns of human language.

In line with these structural distortions, low level patterns of differentiation also emerge. A simple CNN efficiently distinguishes synthetic from natural sequences across domains, indicating that the deviations introduced by generative models are systematically detectable. We noted a complete model collapse of Evo 2 when generation extended beyond approximately 400 kb, limiting the usefulness of the large context length. Even within the initial 300 kb we observed a marked quality drop as the generation as the distance from the seed increases. This is reflected in the performance of our classifier model, which is able to distinguish natural from synthetically generated sequences much more accurately when they are sampled from a longer distance from the initial prompt.

These limitations have important implications for the interpretation and application of gLM-generated genomic sequences. Recent work has demonstrated that Evo and Evo 2 can generate functional synthetic genomes, including viable bacteriophages with desirable biological properties ^26^. However, our findings indicate that functional viability does not guarantee faithful recapitulation of the organizational principles and evolutionary constraints that characterize natural genomes, particularly at large context lengths. Our conclusions are consistent with previous work indicating limitations of synthetic phages generated with megaDNA ^8^. While synthetic sequences hold promise for applications such as phage therapy and personalized therapeutic construct design ^27,28^, their broader biological authenticity remains uncertain; using them to benchmark genomic algorithms, infer evolutionary processes, design regulatory elements in natural genomic contexts, or test hypotheses about genome organization may currently lead to misleading conclusions. Because functional viability and compositional authenticity are decoupled in these synthetic genomes, conclusions about genome biology drawn from gLM-generated sequences must be carefully qualified based on the specific question being addressed.

Our findings argue for the development of more sophisticated genomic architectures tailored specifically to cases where gLMs are used to address biological questions rather than solely for functional sequence generation. Multi-modal approaches that integrate genomic sequences with complementary data modalities, such as transcriptomic, epigenetic, or chromatin accessibility profiles, may better capture the complex organizational principles governing genome architecture ^2,29,30^. Biological questions span a diverse range of applications, from variant effect prediction and regulatory element identification to understanding evolutionary constraints and modeling genome-wide organization patterns. For applications requiring authentic biological fidelity, future genomic foundation models could incorporate explicit evolutionary priors, architectural constraints that account for long-range dependencies and hierarchical genomic organization, and training objectives that reward not only sequence plausibility but also adherence to known biological constraints. Rather than relying solely on token-level sequence prediction, these enhanced architectures could leverage auxiliary biological signals during training to learn representations that better reflect the principles shaping natural genomes. Finally, the quantitative benchmarks introduced here may serve as a foundation for ongoing capability evaluation, enabling the community to track whether and when generative models begin to produce sequences that are indistinguishable from their natural, a question with relevance to both sequence design applications and biosecurity governance.

## Code and data availability

The GitHub code for this project can be found in: https://github.com/Georgakopoulos-Soares-lab/Synthetic_genomes_benchmark

The dataset used in this study is available on Zenodo at: https://doi.org/10.5281/zenodo.18226182

A container with the environment used in this project to facilitate reproducibility through docker/apptainer is provided at: https://doi.org/10.5281/zenodo.15194473

## Supplementary Material

**Supplementary Figure 1:**
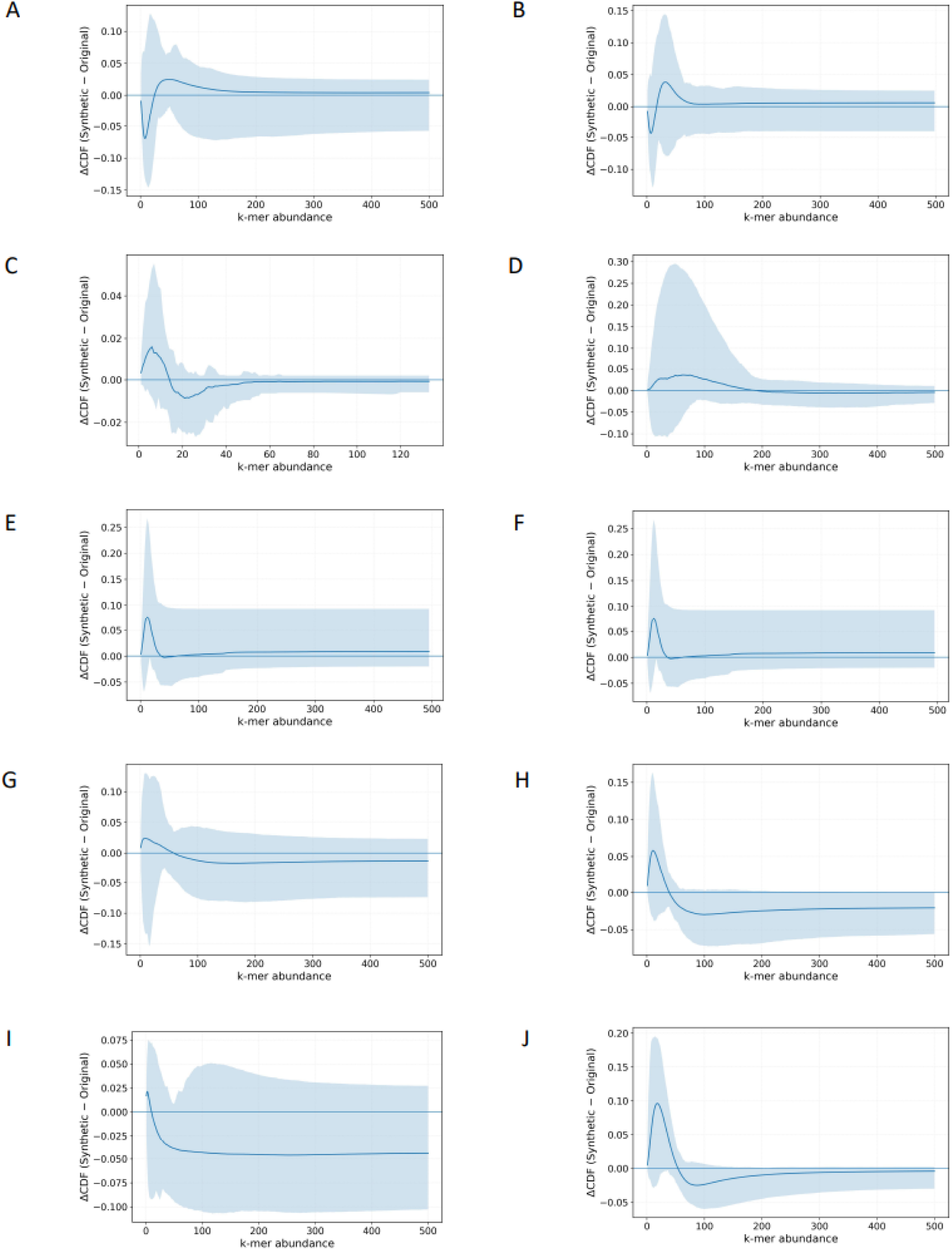
ΔCDF of normalized k-mer abundances (Synthetic − Original) across taxa. Curves show the mean difference in cumulative distributions across genomic windows, with 95% confidence intervals. Deviations from zero indicate redistribution of k-mer frequencies between original and synthetic genomes across abundance levels. Panels A–J correspond to: **(A)** *Homo sapiens*, **(B)** *Mus musculus*, **(C)** Kitrinoviricota, **(D)** Peploviricota, **(E)** Preplasmiviricota, **(F)** Uroviricota, **(G)** Archaea, **(H)** Chlamydiota, **(I)** Mycoplasmatota, and **(J)** Pseudomonadota.

**Supplementary Figure 2:**
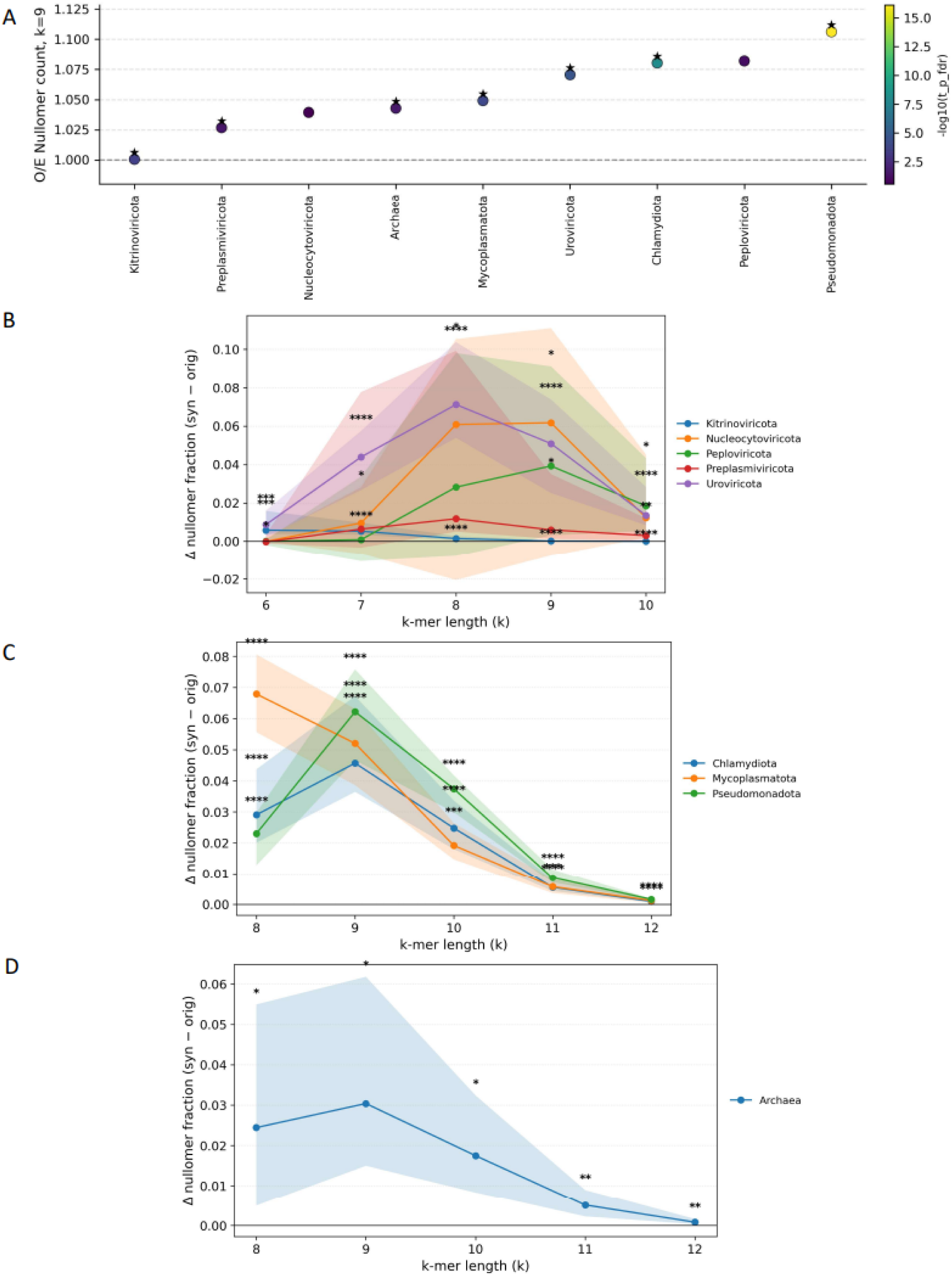
Relative shift in nullomer content in synthetic genomes. **A)** Number of observed (wild-type) and expected (synthetic) nullomers across organismal genomes **(B–D)** Phylum-level nullomer trends showing the median difference in nullomer fraction between synthetic and original genomes (Δ = syn − orig) across species, with 95% bootstrap confidence intervals. **(B)** Viral phyla (k = 6-10). **(C)** Bacterial phyla (k = 8-12). **(D)** Archaea (k = 8-12). Significance indicates deviation from zero based on a Wilcoxon signed-rank test across species, with false discovery rate correction applied separately for viral and bacterial phyla and no correction for Archaea.

**Supplementary Figure 3:**
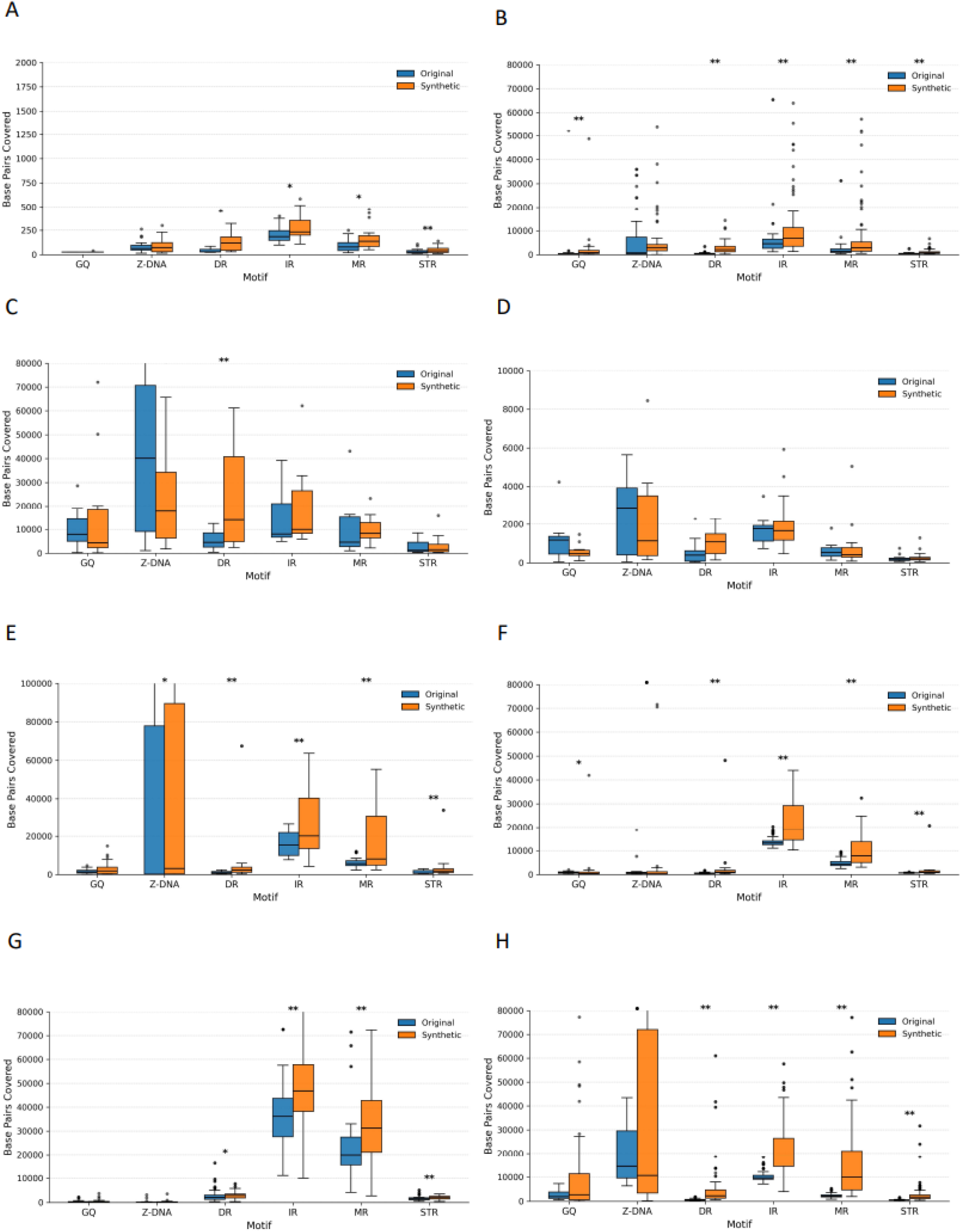
Comparison of non-B DNA motif content in original versus Evo-2-generated synthetic genomic sequences. **A-H.** Boxplots showing the distribution of base-pair coverage for six major non-B DNA motif classes, G-quadruplexes (GQ), Z-DNA, direct repeats (DR), inverted repeats (IR), mirror repeats (MR), and short tandem repeats (STR), in original (blue) and synthetic (orange) genomic windows. Panels correspond to the following species, in order: **A**. *Kitrinoviricota* **B.** *Uroviricota*, **C.** *Peploviricota*, **D.** *Preplasmiviricota*, **E.** *Archaea*, **F.** *Chlamydiota*, **G.** *Mycoplasmatota*, and **H**. *Pseudomonadota*

**Supplementary Table 1: Species used**

[List of species used](https://docs.google.com/spreadsheets/d/17yeRsFoopdcnbC0runwkQo8vJ3JryjmtWZTG58eDGU4/edit?gid=0#gid=0)

**Supplementary Table 2: Species-level divergence between wild-type and synthetic k-mer spectra**

[Species-level divergence between wild-type and synthetic k-mer spectra](https://docs.google.com/spreadsheets/d/1OQvR2Q4FtKkMpvZW2XCnfKrvRnMrVZflNO-BdooiKjQ/edit?gid=0#gid=0)

**Supplementary Table 3: Species-level divergence of frequency chaos game representations between wild-type and synthetic genomes**

[Species-level divergence of frequency chaos game representations between wild-type and synthetic genomes](https://docs.google.com/spreadsheets/d/1nKvNS72BZNiOp0kfApbBvxmPD_CO0NzQrhKuOq7836E/edit?gid=0#gid=0)

**Supplementary Table 4: Nullomers p-value and counts comparison**

[Nullomers p-value and counts comparison](https://docs.google.com/spreadsheets/d/1J3NDRscbwOg9mnNnZ97Fk82HtpUGzPetNmEbtZ3jD2w/edit?gid=0#gid=0)

## References

1. Minaee, S., et al. Large Language Models: A Survey. (2024).

2. Genomic language models: opportunities and challenges. Trends in Genetics 41, 286–302 (2025).

3. Brixi, G. et al. Genome modeling and design across all domains of life with Evo 2. Genomics (2025).

4. Shao, B. & Yan, J. A long-context language model for deciphering and generating bacteriophage genomes. Nature Communications 15, 9392 (2024).

5. Pannu, J. et al. Dual-use capabilities of concern of biological AI models. PLoS Comput. Biol. 21, e1012975 (2025).

6. Sanabria, M., Hirsch, J. & Poetsch, A. R. Distinguishing word identity and sequence context in DNA language models. BMC Bioinformatics 25, 1–12 (2024).

7. Consens, M. E., Li, B., Poetsch, A. R. & Gilbert, S. Genomic language models could transform medicine but not yet. npj Digital Medicine 8, 1–4 (2025).

8. Ratcliff, J. Transformer model generated bacteriophage genomes are compositionally distinct from natural sequences. NAR Genom. Bioinform. 6, lqae129 (2024).

9. Chor, B., Horn, D., Goldman, N., Levy, Y. & Massingham, T. Genomic DNA k-mer spectra: models and modalities. Genome Biology 10, R108 (2009).

10. Jeffrey, H. J. Chaos game representation of gene structure. Nucleic Acids Res 18, 2163–2170 (1990).

11. Acquisti, C., Poste, G., Curtiss, D. & Kumar, S. Nullomers: really a matter of natural selection? PLoS One 2, e1022 (2007).

12. Vergni, D. & Santoni, D. Nullomers and High Order Nullomers in Genomic Sequences. PLoS One 11, e0164540 (2016).

13. Georgakopoulos-Soares, I., Yizhar-Barnea, O., Mouratidis, I., Hemberg, M. & Ahituv, N. Absent from DNA and protein: genomic characterization of nullomers and nullpeptides across functional categories and evolution. Genome Biology 22, 245 (2021).

14. Koulouras, G. & Frith, M. C. Significant non-existence of sequences in genomes and proteomes. Nucleic Acids Res 49, 3139–3155 (2021).

15. Wang, G. & Vasquez, K. M. Dynamic alternative DNA structures in biology and disease. Nat Rev Genet 24, 211–234 (2023).

16. Bochalis, E. et al. Non-B DNA structures and their contributions to genetic diversity, aging, and disease. Nucleic Acids Res 54, (2026).

17. Hsieh, C.-P., et al. RULER: What’s the real context size of your long-context language models? arXiv [cs.CL] (2024).

18. O’Leary, N. A. et al. Reference sequence (RefSeq) database at NCBI: current status, taxonomic expansion, and functional annotation. Nucleic Acids Res 44, D733–45 (2016).

19. Benson, D. A. et al. GenBank. Nucleic Acids Res 42, D32–7 (2014).

20. Kokot, M., Dlugosz, M. & Deorowicz, S. KMC 3: counting and manipulating k-mer statistics. Bioinformatics 33, 2759–2761 (2017).

21. Wang, G. et al. ZSeeker: an optimized algorithm for Z-DNA detection in genomic sequences. Brief Bioinform 26, (2025).

22. Bedrat, A., Lacroix, L. & Mergny, J.-L. Re-evaluation of G-quadruplex propensity with G4Hunter. Nucleic Acids Res 44, 1746–1759 (2016).

23. Cer, R. Z. et al. Non-B DB: a database of predicted non-B DNA-forming motifs in mammalian genomes. Nucleic Acids Res 39, D383–91 (2011).

24. Grant, C. E., Bailey, T. L. & Noble, W. S. FIMO: scanning for occurrences of a given motif. Bioinformatics 27, 1017–1018 (2011).

25. Ovek Baydar, D., et al. JASPAR 2026: expansion of transcription factor binding profiles and integration of deep learning models. Nucleic Acids Res. (2025) doi:10.1093/nar/gkaf1209.

26. King, S. H. et al. Generative design of novel bacteriophages with genome language models. Synthetic Biology (2025).

27. Lenneman, B. R., Fernbach, J., Loessner, M. J., Lu, T. K. & Kilcher, S. Enhancing phage therapy through synthetic biology and genome engineering. Curr Opin Biotechnol 68, 151–159 (2021).

28. Yin, C. et al. Iterative deep learning design of human enhancers exploits condensed sequence grammar to achieve cell-type specificity. Cell Syst 16, 101302 (2025).

29. Consens, M. E. et al. Transformers and genome language models. Nat. Mach. Intell. 7, 346–362 (2025).

30. Javed, N. et al. A multi-modal transformer for cell type-agnostic regulatory predictions. Cell Genom. 5, 100762 (2025).

